# Exploration of Western Himalayan region for identification of gold nanoparticles synthesizing bacteria

**DOI:** 10.1101/164103

**Authors:** Rajni Kant Thakur, Poonam Shirkot

## Abstract

The development of eco-friendly methods for the synthesis of nanomaterial shape and size is an important area of research in the field of nanotechnology. In present study an indigenous bacterial strain GPI-1 has been isolated from a local gold mine Khaltunala. It was characterized morphological, biochemically and also by using 16S rrna gene technology and was identified as *Bacillus flexus* GPI-1, the phylogeny of this bacterial strain was determined using various bioinformatics tools viz BLASTn and MEGA 5.0. To achieve maximum *invitro* gold nanoparticles synthesis various parameters such as pH, incubation temperature, incubation time period and wavelength were optimized as 6.8, 37°C, 36 hrs, 560 nm respectively. Stable and cubical gold nanoparticles (GNPs) formation with 40-45 nm dimensions were synthesized successfully under *invitro* conditions upon exposure of gold chloride trihydrates (HAuCl_4_) solution to the supernatant of *Bacillus flexus* strain GPI-1. These gold nanoparticles have been characterized by Transmission electron microscope, Fourier transform infrared spectroscopy (FTIR). Thus in the present study successful biosynthesis method of stable and cubical gold nanoparticles in the size range of 12-30 nm using this efficient *Bacillus flexus* GPI-1 strain. Thus in the present study leading to development of an easy bioprocess for synthesis of GNPs of desired size and shape, has been reported and this green route of biosynthesis of GNPs is a simple, economically viable and an eco-friendly process. The use of gold nanoparticles in biomedical research like X-ray computed tomography and magnetic resonance imaging, cancer research, drug delivery applications.

## Introduction

Nanotechnology is a group of emerging technologies in which the structure of matter is controlled at the nanometer scale to produce novel materials and devices that have useful and unique properties. In order to achieve the desired control, a special non-random eutectic environment needs to be available. Although chemical and physical methods may successfully produce pure, well defined nanoparticles, these are quite expensive and potentially dangerous to the environment. As an alternative to toxic and expensive physical methods for nanoparticles fabrication, using microorganisms, plants and algae possess significant potential to synthesize the materials in the nano range and in addition, the toxicity of the by-product would be lesser than the other synthetic methods [1-3]. Physical methods usually require high temperature, vacuum and expensive equipment which makes these techniques uneconomical, whereas chemical methods need different types of toxic and expensive chemicals making nanoparticles production very expensive and hazardous. These disadvantages of both physical and chemical methods for synthesis of nanoparticles greatly limit their applications in various fields and therefore development of reliable, nontoxic and eco-friendly methods for synthesis of nanoparticles are of utmost importance to expand their applications. One of alternatives to achieve this goal is to synthesize the nanoparticles by microorganisms, which leads to another new branch of bionanotechnology. The growing momentum of green nanotechnology, the synthesis of NPs utilizing biological materials could be an improved alternative to toxic chemicals and the expensive physical methods. In general, biological organisms, such as bacteria and fungi [4], plants [5], and algae [6] has been reported for development of eco-friendly and cost-effective approaches [7]. Biosynthesis of nanomaterials has received a significant attention in recent times owing to the use of mild experimental conditions such as temperature and pH. These microorganisms play an important role in remediation of metals through reduction of metal ions and some of these microorganisms can survive and grow even at high metal ion concentrations. These are often exposed to extreme environmental conditions, forcing them to resort to specific defence mechanisms to quell such stresses, including the toxicity of foreign metal ions or metals [8]. The biological agents secrete a large amount of enzymes, which are capable of hydrolyzing metals and thus bring about enzymatic reduction of metals ions [9]. The biomass used for the synthesis of nanoparticles is simpler to handle, gets easily disposed of in the environment and also the downstream processing of the biomass is much easier. The mechanism of extracellular synthesis of nanoparticles using microbes is basically found to be nitrate reductase-mediated synthesis [10]. This enzyme nitrate reductase helps in the bioreduction of metal ions and synthesis of nanoparticles. A number of researchers supported nitrate reductase for extracellular synthesis of nanoparticles [11-15]. One of the chief applications of nanotechnology is the use of gold nanoparticles in biomedical research like X-ray computed tomography and magnetic resonance imaging [16], cancer research [17], drug delivery applications [18], and its optical properties for cancer diagnosis and photo thermal therapy. In the present study, an indigenous gold nanoparticle synthesizing bacteria has been isolated from biofilm samples of a local goldmine. This bacterium has been identified as *Bacillus flexus* strain GPI-1 after morphological, biochemical and molecular characterization by 16S *rrna* gene technology. Extracellular, spherical, monodispersed/small clusters of gold nanoparticles were successfully produced. It is an eco-friendly method for the bacteria mediated synthesis of gold nanoparticles by the reduction of HAuCl_4_ ions using the broth of *Bacillus flexus* GPI-1 is reported and the bioreduction process was monitored by the UV-visible spectroscopy at 560 nm. The nanostructure and size of the synthesized gold nanoparticles were characterized by transmission electron microscopy (TEM) and Fourier transformation infrared spectroscopy (FTIR) was used to understand the biomolecules responsible for the biosynthesis.

## Material methods

### Isolation and identification of gold nanoparticles and bacteria

A survey was conducted for selection of various sites of goldmine at Khaltunala, in Solan district of Himachal Pradesh. Forty-three samples in form of water, soil, biofilm, pebbles, stalagmite and rock matting were collected and stored at 4°C till further experimentation. Three different culture media *viz.*, Nutrient agar medium, Eosin methylene blue agar medium and Luria bertani medium were investigated for isolation of gold nanoparticles synthesizing bacterial isolates. One gram of soil, pebbles and stalagmite /1.0 ml water and biofilm samples were used for isolation of bacterial isolates through using serial dilution technique. Each of the morphologically distinctive colonies was transferred to fresh broth medium and incubated at 37°C for 24 hrs. Turbid cultures were streaked on plates of solidified growth medium. Individual colonies were restreaked repeatedly, and the axenic cultures thus obtained were stored at 4°C.

### Morphological characterization

All the bacterial isolates obtained in previous step were further studied for various morphological characters. Various morphological colony descriptors of colour, size, optical property and elevation of the colonies and various microscopic characteristics studied were gram reaction, shape and arrangement of cells and spore formation.

**Quantitative screening of bacterial isolates for gold nanoparticles synthesis ability** Assessment of all forty-three bacterial isolates for their ability to synthesize gold nanoparticles was carried out, transferring 1 % of the inoculum (overnight culture) of each bacterial isolate into 50 ml nutrient broth followed by incubation at 37°C for 24-48 hrs at 150 rpm. Supernatant of each bacterial culture was collected by centrifugation at 8500 rpm for 15 minutes at 4°C. Ten ml of each supernatant was mixed with 10 ml of 1mM solution of (HAuCl_4_) and incubated at 37°C for 240 hrs. Formation of gold nanoparticles was monitored from 0-240 hrs, with an interval of 12 hrs, which was confirmed by colour change of the solution from light yellow to red wine/purple colour. This formation of gold nanoparticles was also confirmed using spectrophotometer (Spectronic 20, Milton Roy Company) at two different wavelengths of 540 and 560 nm.

### Biochemical characterization and molecular characterization

Eleven bacterial isolates selected were investigated further using various biochemical characters *viz* beta galactosidase, hydrolysis oxidase, hydrolysis of starch, citrate utilization D-glucose, D-mannitol, fructose, galactose, lactose, maltose, indole production, Voges-Proskauer reaction, D-mannose, rhamnose using standard assays. DNA extraction from selected bacterial isolates was carried out using genomic DNA extraction Mini kit (Real Genomics). Presence of DNA and its quality was checked using 1.0% agarose gel and was viewed by UV trans-illuminator. The DNA of GPI-1 strain was selectively amplified using 16S rrna gene PCR technology. Universal primers B27F (5`-AGAGTTTGATCCTGGCTCAG-3’U1492R) and (5`-GGTTACCTTGTTACGACTT-3’) for 16S *rrna* gene were used for the experiment and the eluted and purified DNA of GPI-1 was sequenced. The sequence has been submitted to NCBI with accession number KP219454. To gain insight of the evolutionary pattern, phylogenetic tree was constructed using MEGA 5.0 bioinformatics tool. Neighbour-Joining (NJ) technique of mathematical averages (UPGMA) was used. Nanoparticles obtained were analysed using various techniques such as, Fourier transform infrared spectroscopy (FTIR and Transmission electron microscope.

### Optimization of culture conditions for maximum gold nanoparticles synthesis by selected bacterial isolate

The culture conditions for useful and prized microorganisms are generally optimized to obtain higher yields of their useful products. In the present study the selected bacterial isolate strain GPI-1 was investigated to study the effect of different parameters of culture conditions such as incubation time, temperature, pH and wavelength for maximum gold nanoparticles synthesis.

### Effect of incubation time

Effects of different incubation times were ranging from 0-72 hrs for maximum gold nanoparticles synthesis investigated, and the optimum incubation time leading to maximum gold nanoparticles production of gold nanoparticles was selected for further experiments.

### Effect of incubation temperature

Effect of incubation temperature for maximum gold nanoparticles synthesis was studied at a temperature range of 10-50°C using nutrient broth optimum incubation time. The optimum temperature leading to maximum gold nanoparticles production of gold nanoparticles was selected for further experiments.

### Effect of pH

Assessment of optimum pH for maximum gold nanoparticles synthesis by selected bacterial isolate, and pH range of 5.0-7.5 was examined using nutrient broth medium at an optimum temperature and optimum time. The best condition leading to maximum gold nanoparticles production of gold nanoparticles was selected for further experiments.

### Effect of wavelength

Effects of different wavelength for the maximum values of gold nanoparticles synthesis were assessed ranging from 400-650 nm. The optimum wavelength leading to maximum readings of gold nanoparticles synthesis was selected for further experiments.

### Fourier Transform Infrared Spectroscopy

This technique was used to study different biomolecules which were involved in biosynthesis of gold nanoparticles. Microcup was washed with 100% absolute ethanol. 10 ul sample was filled in a 2 mm internal diameter microcup and loaded onto the FTIR set at 26°C±1°C. The samples were scanned in the range of 4,000 to 400 cm^-1^ using a Fourier transform infrared spectrometer (Thermo Nicolet Model 6700, Waltham, MA, USA). The spectral data obtained were compared with the reference chart to identify the functional groups present in the sample.

### Transmission electron microscope

TEM studies were carried out to study different shapes and sizes of nanoparticles,using Jeol 2100 microscope operating at 120 kV accelerating voltage. Samples were prepared by placing a drop of *invitro* synthesis gold nanoparticles solutions on carbon-coated TEM grids. The films on the TEM grids were allowed to dry for 5 min at room temperature before analysis.

## Results and Discussion

Pure colonies were isolated from pebble sample were characterized for their morphological and physiological characteristics by various biochemical tests using the Bergeys Manual of Determinative Bacteriology [20] as summarized in (Table 1). *Bacillus flexus* GPI-1 was found to be gram positive, spore forming, motile, rod shaped and aerobic bacteria. The isolated strain GPI-1 gave positive results for beta galactosidase, hydrolysis oxidase, hydrolysis of starch, citrate utilization and acid production from D-glucose, D-mannitol, fructose, galactose, lactose, maltose. The strain gave negative results indole production, Voges-Proskauer reaction, no acid production from D-mannose, rhamnose (Table1). Molecular characterization was carried out using 16S r DNA-PCR technology. Total genomic DNA of the GPI-1 bacterial isolate was extracted successfully using genomic DNA extraction Mini kit (Real Genomics) and was selectively amplified with universal primers for 16S *rrna* gene followed by agarose gel electrophosis leading to a single clear band. Which was eluted, purified and sequenced. The sequence was submitted to NCBI with accession number Genbank KP219454. BLASTn analysis depicted homology of GPI-1 bacterial isolate with other *Bacillus* species. To gain insight into evolutionary pattern, phylogenetic tree was constructed using MEGA 5.0 bioinformatic tool [21]. The bootstrap analysis values identified the bacterial isolate GPI-1 as *Bacillus flexus* (Figure 1), Multiple sequence alignment of query nucleotide sequence of maximum gold nanoparticles synthesizing indigenous *Bacillus flexus* strain GPI-1 was performed with that of the selected nucleotide sequences using ClustalW program and pairwise percent similarity score of these selected fifteen nucleotide sequences obtained from NCBI database with test isolate GPI-1 from goldmine, elucidates that sequence-1. *Bacillus flexus* strain GPI-1 showed maximum similarity score of 99% with *Bacterium* BW4SW28 (Figure 1).

**Table: 1.**
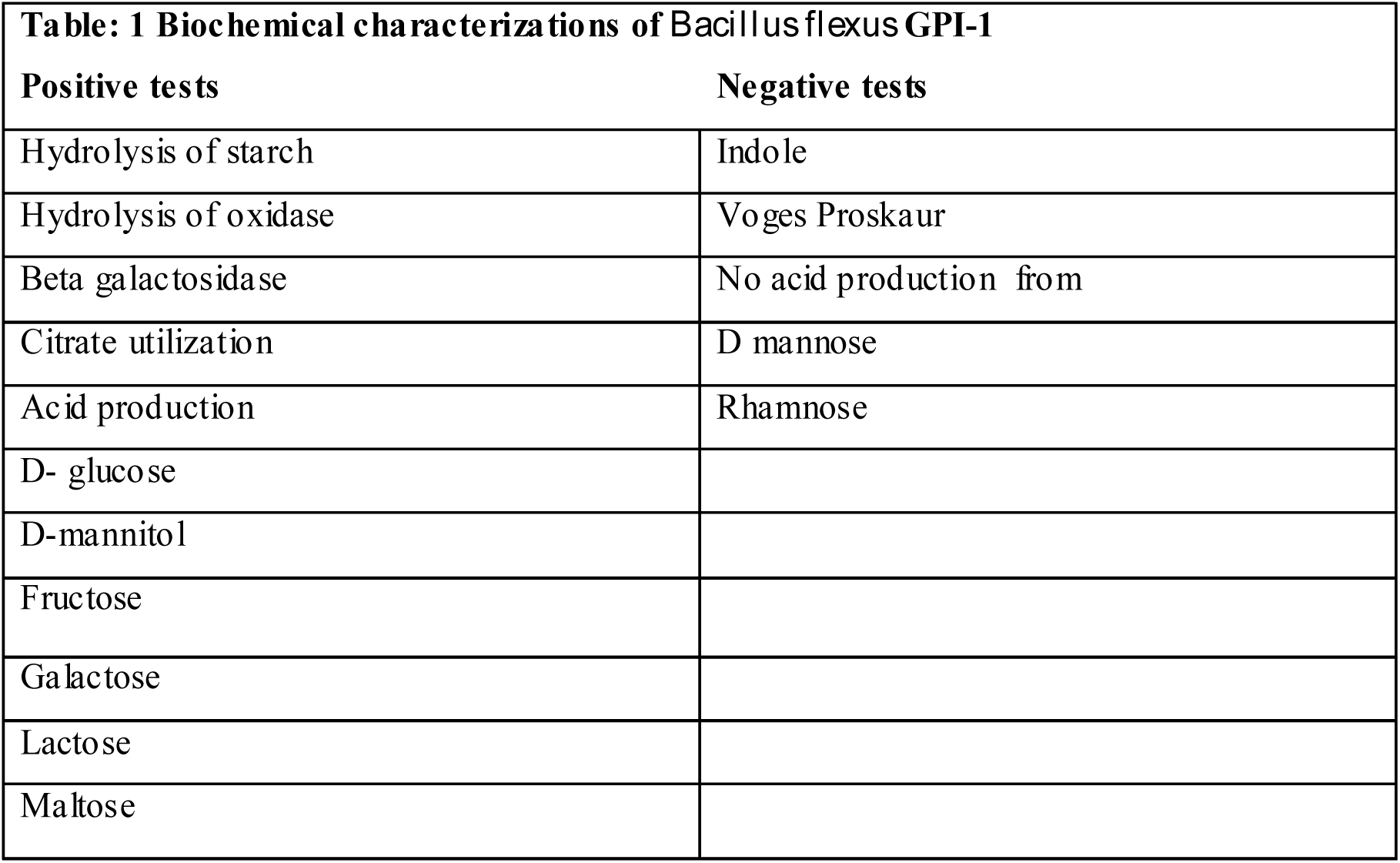
Biochemical characterizations of *Bacillus flexus* GPI-1

**Figure 1.**
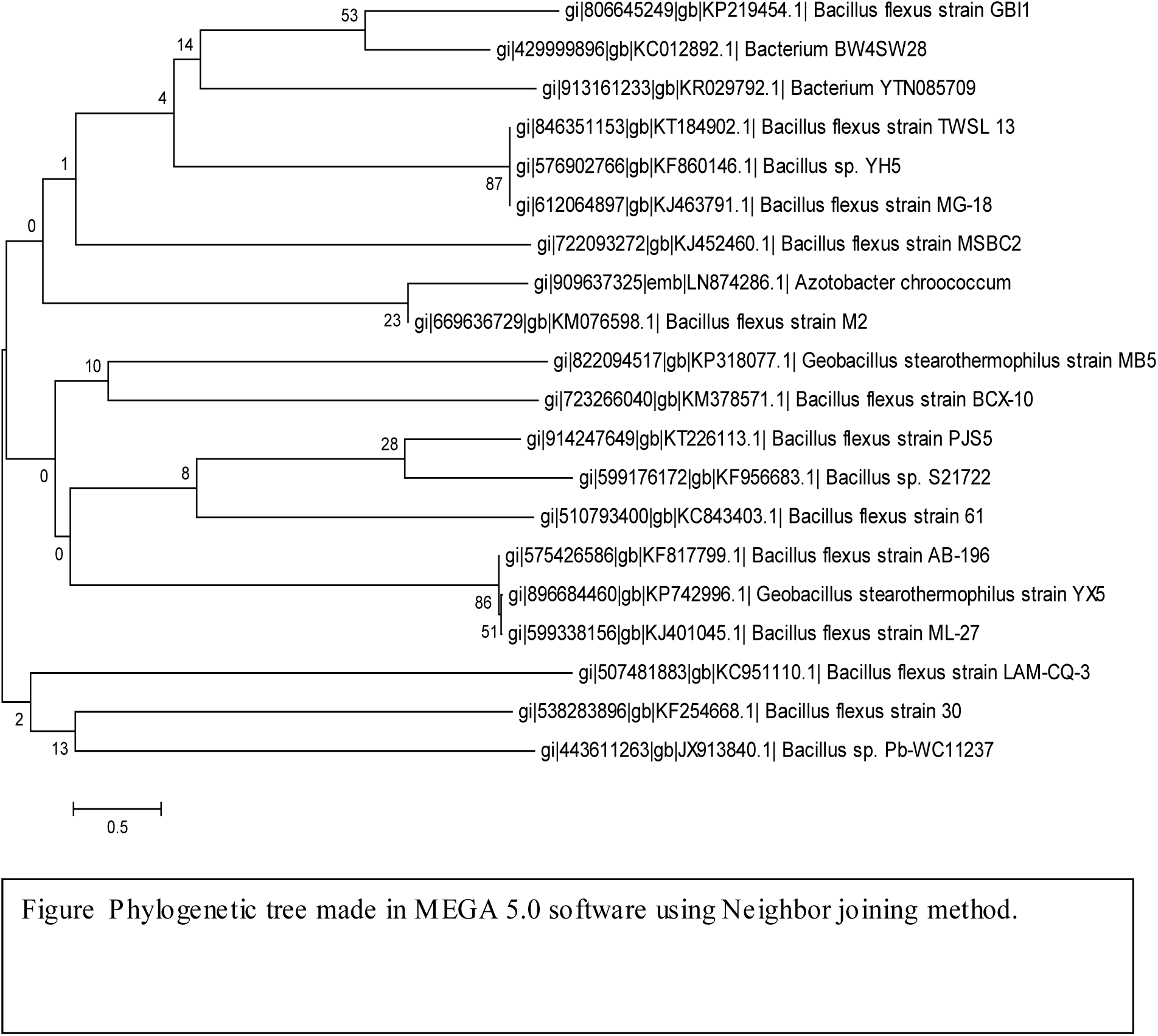
Phylogenetic tree made in MEGA 5.0 software using Neighbor joining method.

### Characterization of gold nanoparticles synthesized by Bacillus flexus GPI-1

Addition of gold chloride solution (HAuCl_4_) added into the supernatant of *Bacillus flexus* GPI-1 changed the colour progressively from light yellow to red at a temperature 37°C of predicting formation of gold nanoparticles. The kinetics of reaction was further studied using the spectronic 20 D by recording spectra from colloidal gold solution obtained after, mixing 10 ml of 1mM gold chloride solution with 10 ml of supernatant of *Bacillus flexus* GPI-1. The spectra revealed a strong absorption at wavelength of 560 nm and it increased upto 36 hrs and then decreased. After 36 hrs of incubation, as synthesis of GNPs initiated only after 6hrs of incubation and showed the presence of GNPs in colloidal solution upto 240hrs at 37°C (figure 2a,b). Size of GNPs was observed through transmission electron microscope and images demonstrating gold nanoparticles possessing the average diameter of 45nm as depicted in (figure 3). The fourier transform infrared (FTIR) spectra obtained from the gold chloride solution after interaction with supernatant of *Bacillus flexus* GPI-1 for 24hrs. The spectra showed the occurrence of three bands at 3500cm^-1^, 2100 cm^-1^ 1980 cm^-1^ and 1480 cm^-1^. Peak 3500 cm-^1^ indicates the presence of carboxylic acids, and N-H primary amines. Peak 2100 cm^-1^ refers to C≡C terminal alkynes, peak 1980 cm^-1^ showed the presence of C-N bond, R-N=C=S is specific type of bond, 1480 cm^-1^ peak indicates the presence of C-H bond and confirm that alkyl group is present (figure 4).

**Figure 2a.**
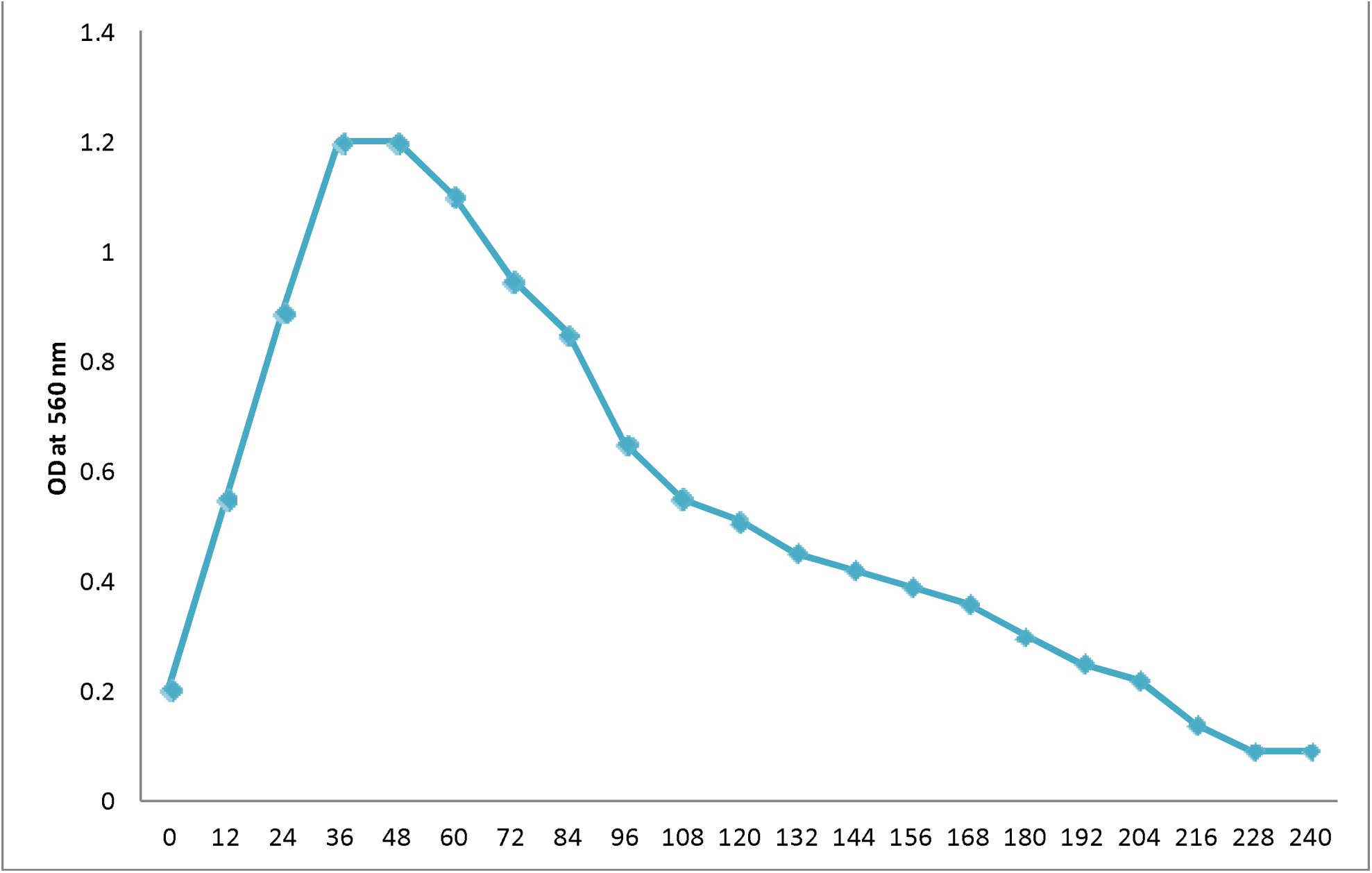
UV-Vis absorption spectra of gold nanoparticles after incubation of supernatant of *Bacillus flexus* GPI-1 with 1mM sodium chloride for time period (0-240 hrs) at 6.8 pH

**Figure 3.**
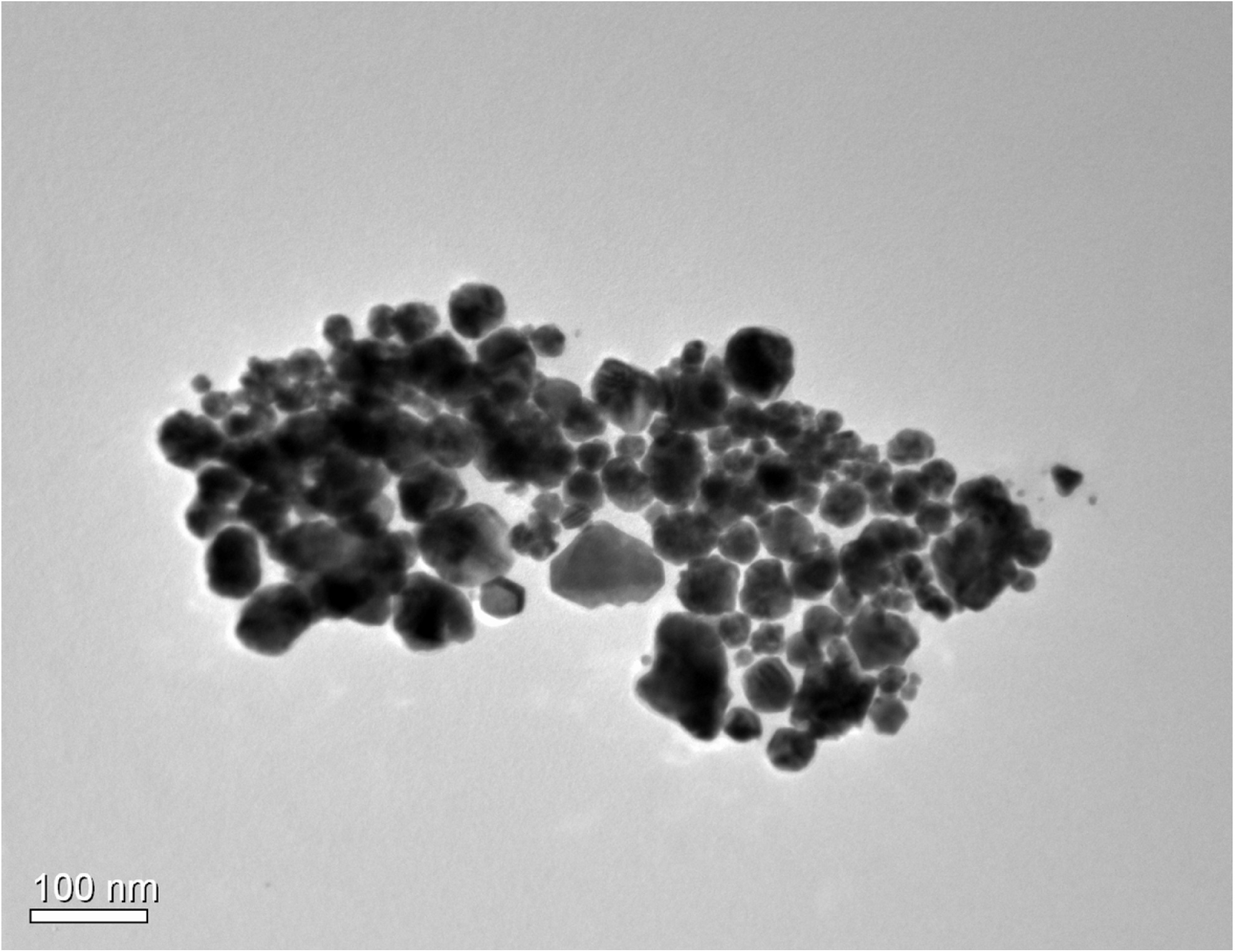
Transmission electron microscope images of indigenous Bacillus flexus GPI-1, showing the different shapes and sizes of nanoparticles.

**Figure 4.**
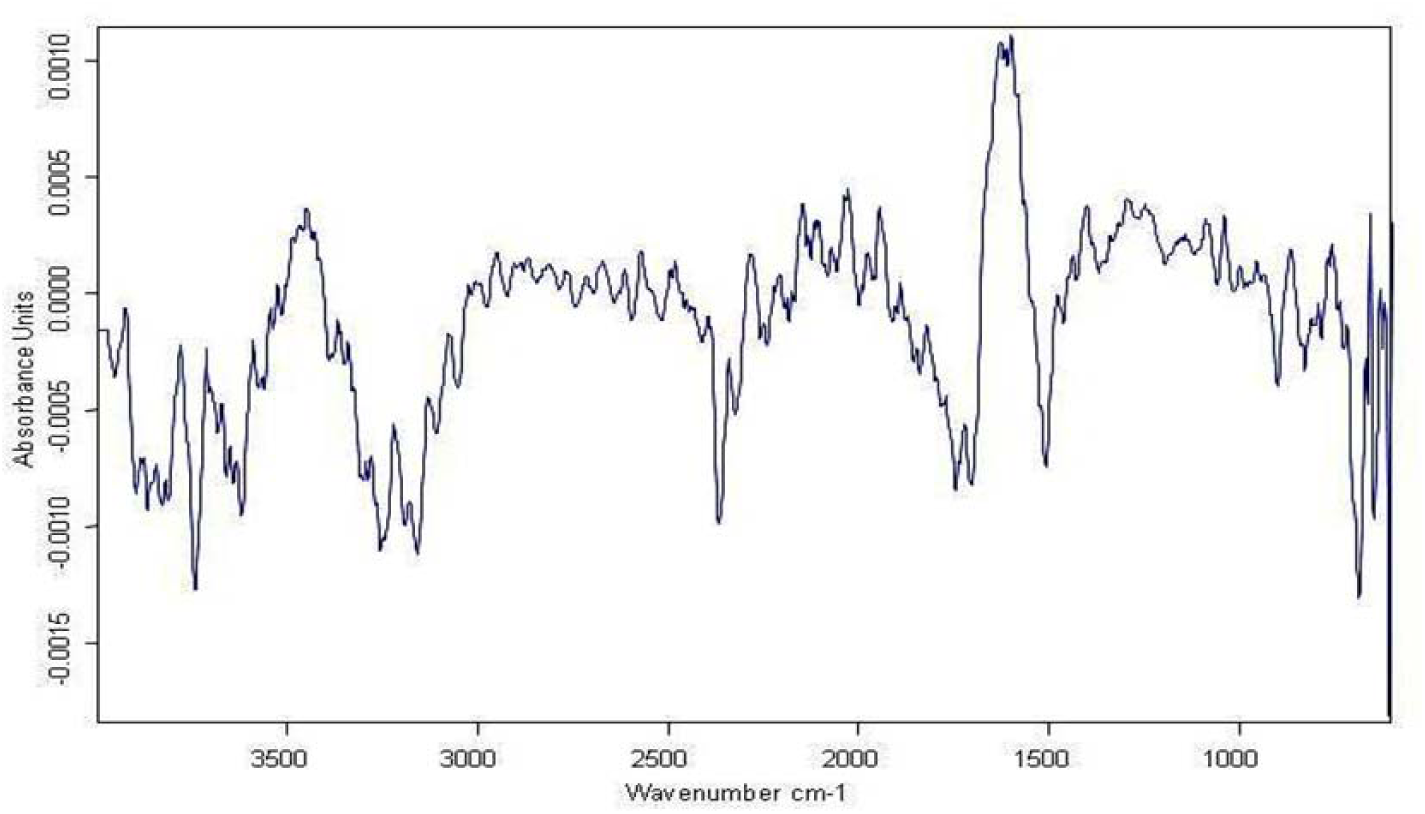
Typical FTIR spectrum of gold nanoparticles generated by *Bacillus flexus* GPI-1

### *In vitro* synthesis of gold nanoparticles by indigenous *Bacillus flexus* strain GPI-1

Extracellular biosynthesis of gold nanoparticles was carried out using supernatant of *Bacillus flexus* strain GPI-1, treated with 1mM gold chloride solution and incubated at 37°C for a time period of 0-240 hrs. Biosynthesis absorption spectra of gold nanoparticles which was indicated by colour change of solution from yellow to red wine (Figure 5) and was further confirmed spectrophotometrically. UV-VIS absorption spectra and the time of incubation course and increase in formation of gold nanoparticles took place upto 36 hrs and remained stable upto 48 hrs and then the values declined upto 240 hrs. Gold nanoparticles formation clearly revealed the gold nanoparticles formation initiated after 6 hrs and studies at two different wavelengths of 540 nm and 560 nm (Figure 2a, b). On the basis of data obtained at 540,560 nm it was quite clear that results obtained at 560nm had a supremacy over 540 nm results, so 560 nm wavelength was selected for further experiments. Gold nanoparticles O.D value sharply increases 6 to 36 hrs and then it showed stability upto 48 hrs.

**Figure 5.**
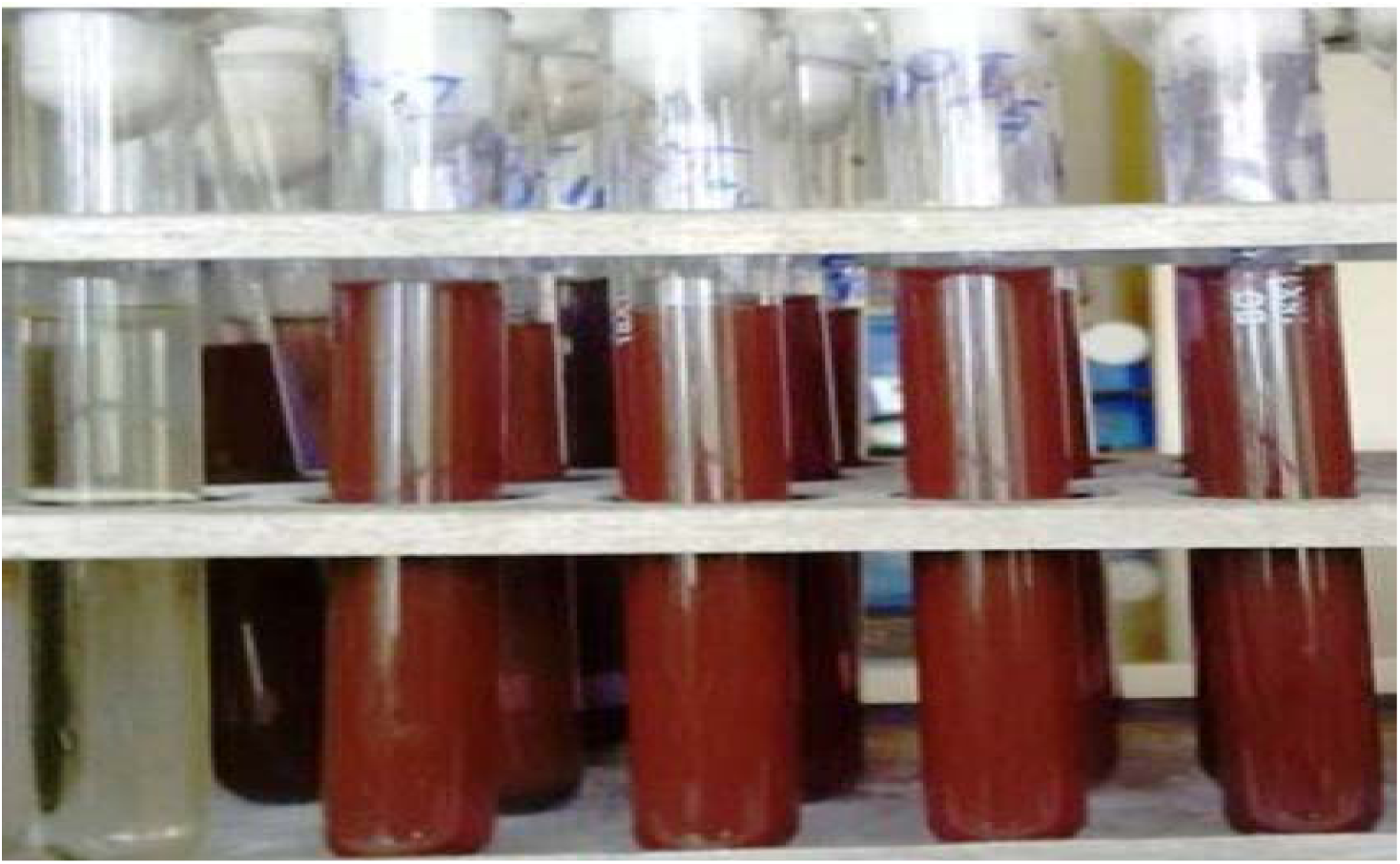
Synthesis of gold nanoparticlese by Bacillus flexus GPI-1

### Optimization of culture conditions for maximum gold nanoparticles synthesis by *Bacillus flexus* strain GPI-1

Traditional optimization has been carried out by monitoring the effect of one factor at a time on an experimental response. While only one parameter is changed, others are kept at a constant level. The bacterial isolate *Bacillus flexus* GPI-1 depicting maximum gold nanoparticles synthesis activity was further optimized to study the effect of different factors such as incubation time, temperature, pH and wavelength on gold nanoparticles synthesis. Effect of pH on biosynthesis of gold nanoparticles by *Bacillus flexus* GPI-1 bacterial supernatant of was studied for 0-72 hrs using 1mM gold chloride solution at a pH range of 5.0, 6.0, 6.8, 7.5, 8.0 (Figure 6a) and it was observed that maximum gold nanoparticles took place at pH: 6.8. Thus pH has been found to be an important parameter affecting gold nanoparticles synthesis [22]. Variation in pH during exposure to gold ions had an impact on the size, shape and number of particles produced per cell [23]. Gold nanoparticles formed at pH 6.8 were predominantly triangles, spherical, hexagons, circular in shape. Whenever pH increases, more competition occurs between protons and metal ions for negatively charged binding sites. Effects of different incubation times for maximum gold nanoparticles synthesis were investigated from 0-72 hrs. The optimum incubation time of 36 hrs, leading to maximum gold nanoparticles production was observed (Figure 6b). Effect of incubation temperature for maximum gold nanoparticles synthesis was studied at a temperature range of 10-50°C using nutrient broth and optimum temperature of 37°C leading to maximum gold nanoparticles synthesis was observed (Figure 6c). It has also been reported that incubation time for maximum gold nanoparticles formation ranges from 30 to 37°C [24]. Effect of different wavelengths for the maximum optical density values of gold nanoparticles synthesis was investigated in the range of 400-650 nm and an optimum wavelength of 560 nm was found to be leading to maximum values of optical density of gold nanoparticles synthesis (Figure 6d).

**Figure 6a.**
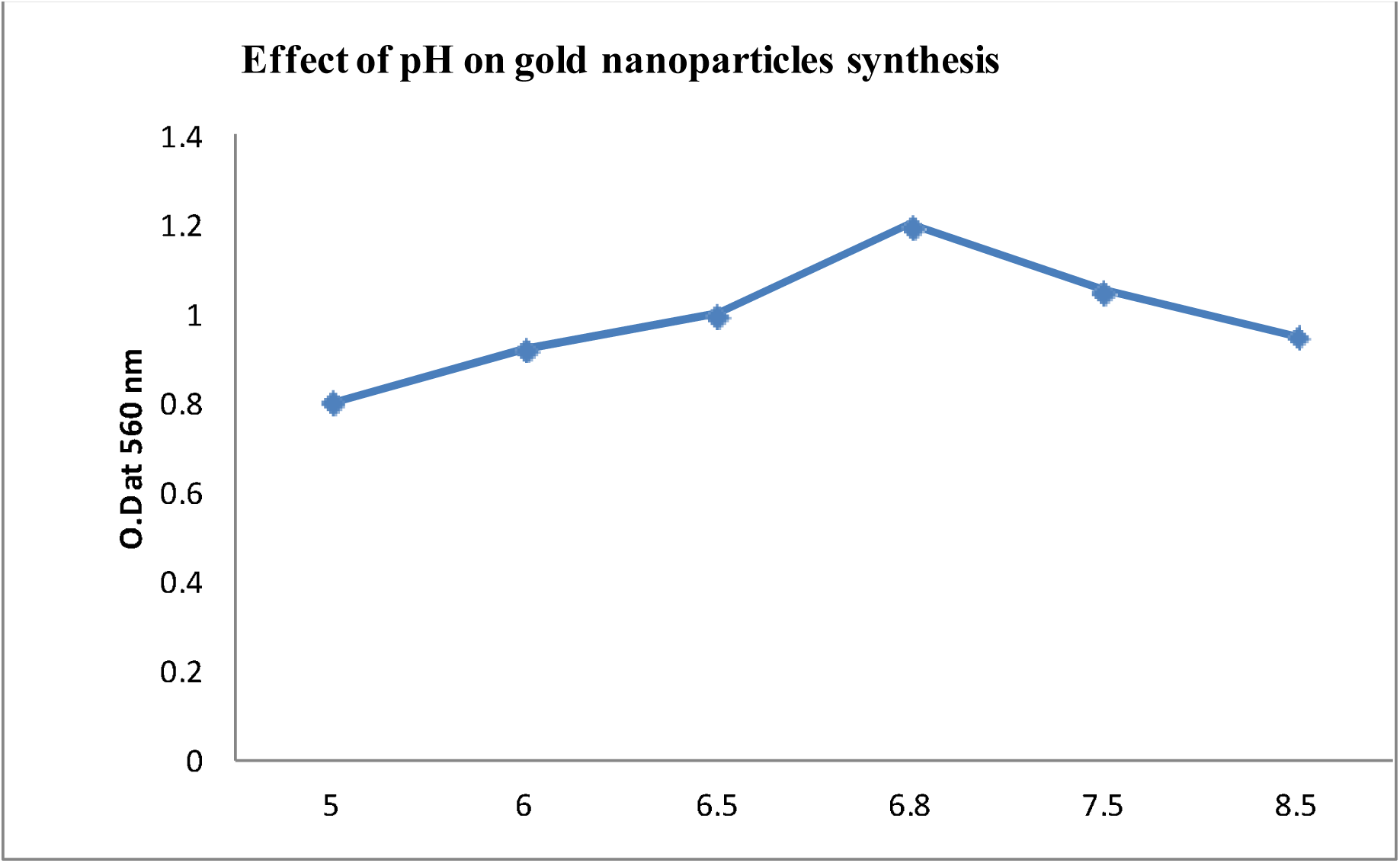
Effect of pH on gold nanoparticles synthesis, and it was found that at 6.8 pH maximum GNPs synthesis take place.

**Figure 6b.**
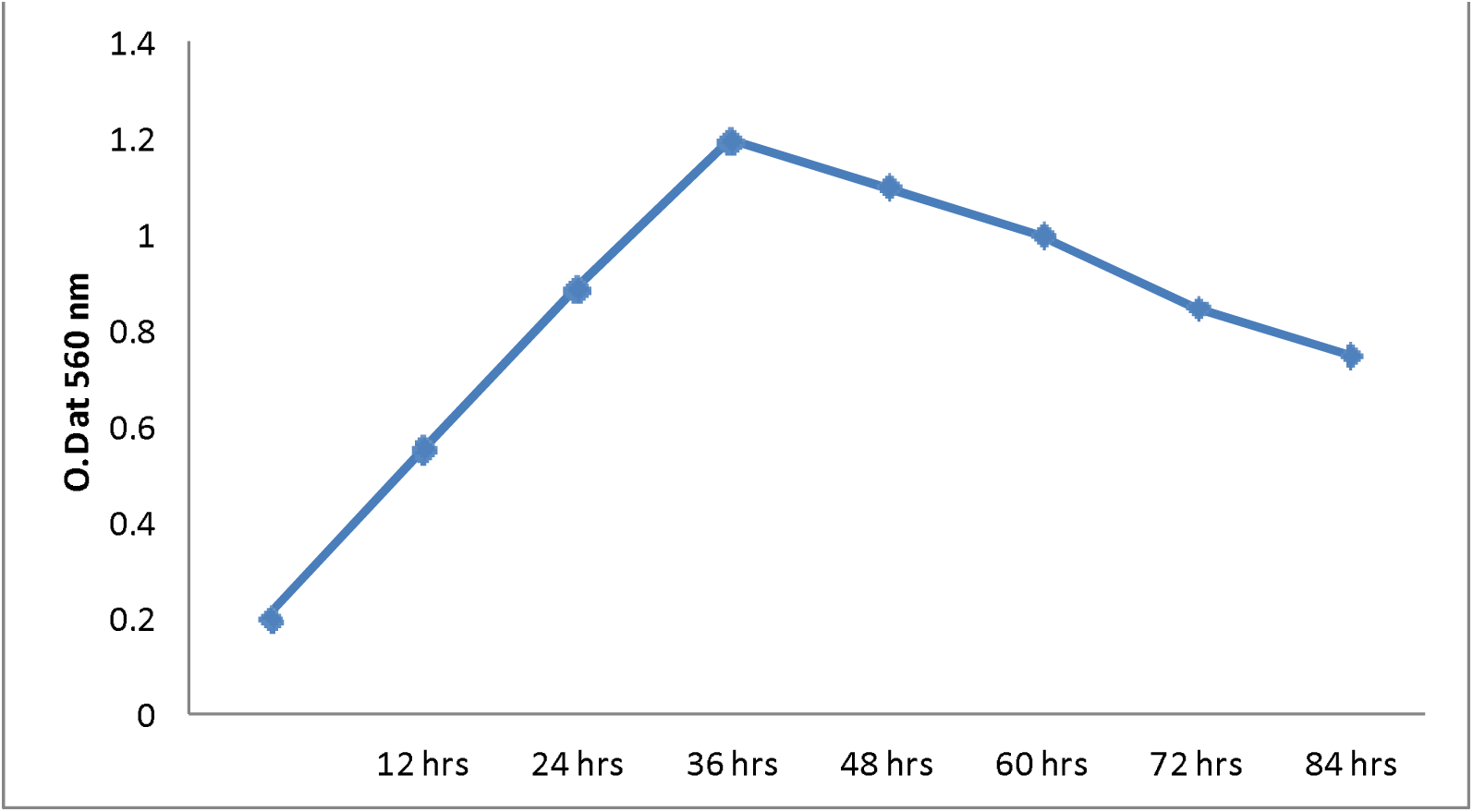
Effect of different wavelengths on measurement of gold nanoparticles.

**Figure 6c.**
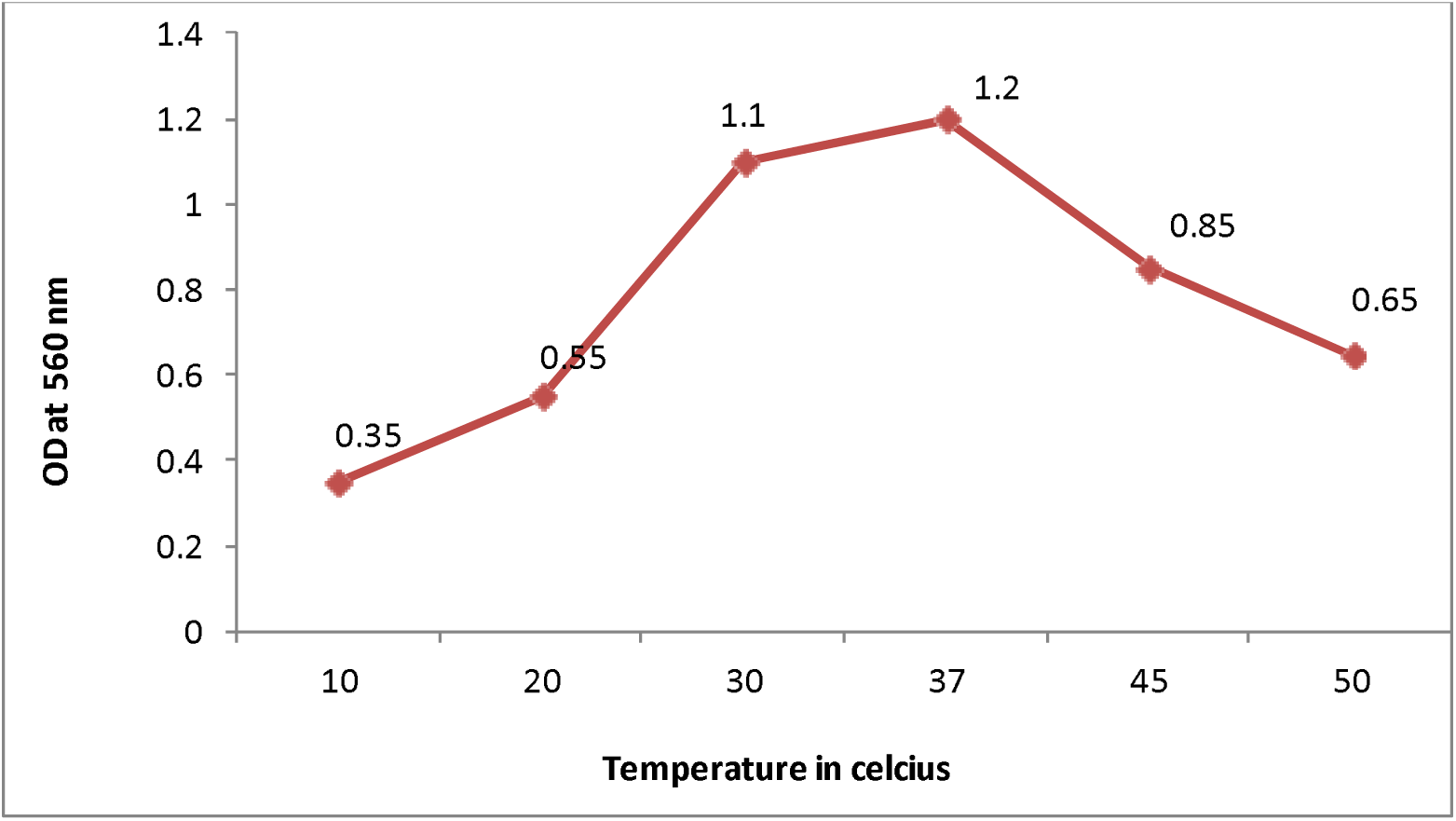
Effect of incubation temperature on gold nanoparticles synthesis.

**Figure 6d.**
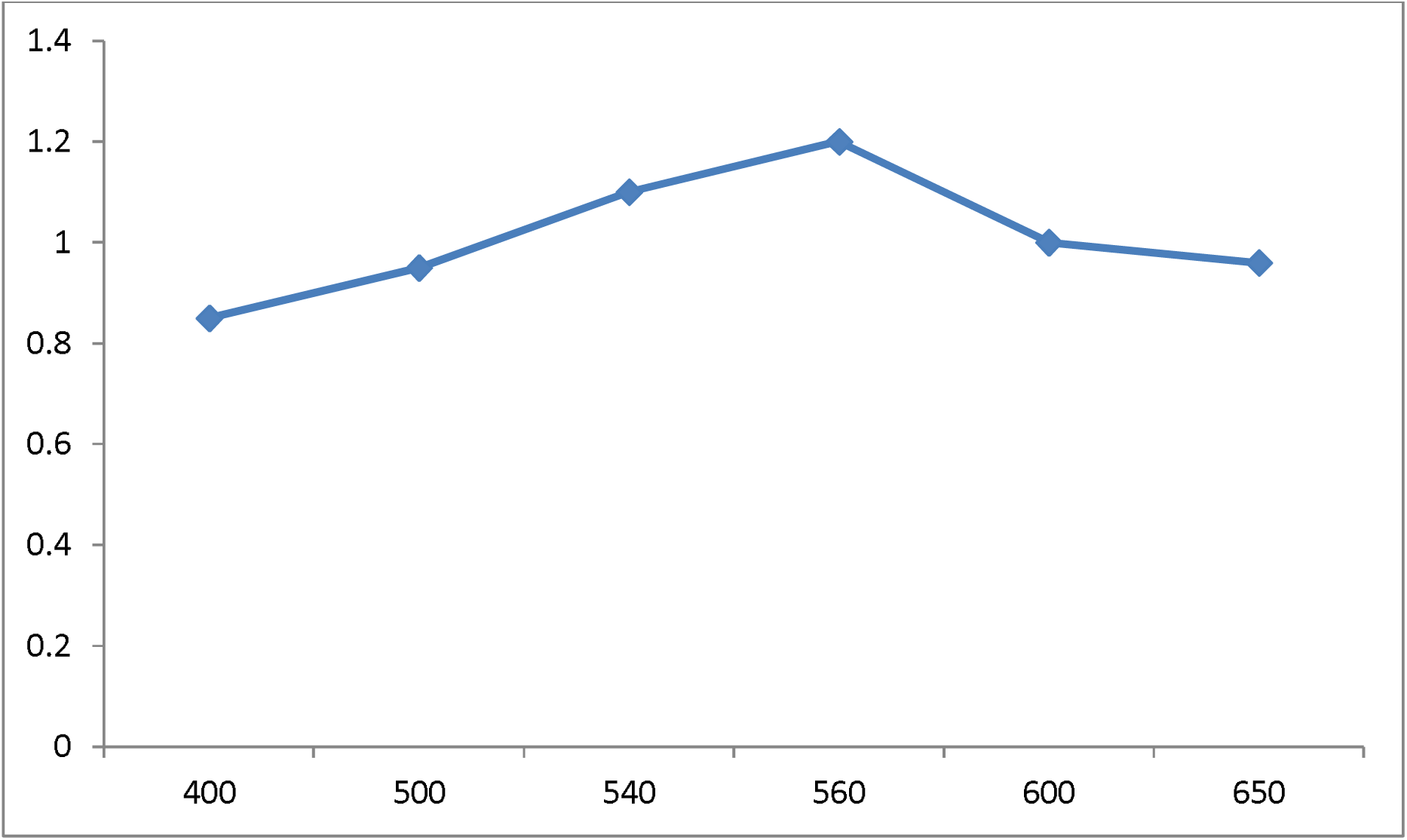
Effect of incubation time on gold nanoparticles synthesis.

## Conclusion

Formation of gold nanoparticles was apparent from the gradual changes in the color of incubated solution from pale yellow to dark purple, but the color of control remained practically unchanged during the entire incubation period. This color changes from pale yellow to dark purple is attributed to surface plasmon resonance. The FTIR measurements were carried out to identify the possible bio-molecules responsible for capping and efficient stabilization of the metal nanoparticles synthesized by *Bacillus flexus*. The spectrum represents different functional groups of adsorbed. It seems that the FTIR spectrum reveal the presence of different functional groups like there may be presence of carboxylic acids, and N-H primary amines, alkyl group helps in synthesized gold nanoparticles. It is believed that the enzyme nitrate reductase could be responsible for bioreduction of metal ions and synthesis of nanoparticles. The bioreduction of gold ions was found to be initiated by the electron transfer from the NADPH-dependent reductase as a electron carrier. Next, the gold ions (Au3^+^) obtain electrons and are reduced to elemental gold (Au^0^) nanoparticles. The secreted proteins and enzymes in the medium are responsibles not only for synthesis of metallic nanoparticles, but also for stabilization of nanoparticles against aggregation. It has been reported first time that the use of *Bacillus flexus* could be used for extracellular synthesis for gold nanoparticles. The stability of the nanoparticles solution could be due to secretion of certain reducing enzymes and capping proteins by the bacterium. Extracellular formation of gold nanoparticles would be advantageous from process point of view, since it would eliminate the need to recover the particles formed within the cells.

## Authors’ contributions

PS conceptualized and designed experiments and provided technical support and helped in drafting the manuscript. RKT conducted experiments, isolation of bacteria, morphological and molecular characterization, invitro synthesis of gold nanoparticles and characterization using FTIR and TEM and helped in drafting the manuscript both author read and approved the final manuscript.

## Author information

RKT is a PhD research scholar currently and member of six international and national science societies. PS is a professor of Biotechnology with 30 years’ research experience and has published research papers in the journals of international and national repute.

## Acknowledgement

The authors thank to Dr. C Raman Suri and Dr. Manoj Raje for the technical guidance and Dr. CK Shirkot and Dr. Garish Sahani for their cooperation.

